# De novo design of metalloenzymes using Metal Mers

**DOI:** 10.64898/2026.10.13.758590

**Authors:** Chuan Shan, Zhuoyun Ren, Jinhao Yu, Yuan Gao, Xiaofang Zhong, Lei Lu

## Abstract

De novo metalloenzyme design for new-to-nature transformations remains challenging, with limited substrate scope and reaction diversity and a need for extensive experimental optimization. Here, we present a strategy based on Metal Mer, a structural unit that maps protein backbone to statistically preferred positions of bound metals, enabling precise metal coordination and efficient metalloenzymes design. Using this strategy, we designed zinc-dependent hydride-transfer enzymes (17/60 active), achieving >99% yield and >99.5:0.5 enantiomeric ratio (e.r.). Redesigned variants catalyzed steroid transformations with >19:1 site- and diastereoselectivity. Furthermore, designed iron enzymes catalyze alkene trifluoromethylazidation (17/36 active), and a single active-site mutation yielded a variant achieving up to 89% yield and 97:3 e.r., rivaling extensively evolved enzymes. Crystal structures closely matched computational models (Cα RMSD <0.75 Å), supporting the accuracy and generality of this approach.

**One Sentence Summary:** We present Metal Mer, a protein structure unit that enables the de novo creation of highly efficient and selective metalloenzymes for new-to-nature reactions without directed evolution.

## Introduction

Biocatalysis provides sustainable access to chiral, densely functionalized building blocks for pharmaceuticals and materials. Metalloenzymes are of particular interest, since metal centers greatly broaden the transformations accessible to organic synthesis. Efforts in this area have primarily focused on introducing non-native metal ions or metal–organic cofactors into natural protein scaffolds, or repurposing native enzymes through directed evolution to enable new-to-nature reactions (*1, 2*). Beyond engineering natural proteins, de novo design builds biocatalysts from first principles rather than from existing scaffolds. In particular, de novo metalloproteins offer the potential to combine the intrinsic reactivity of metal centers with the selectivity, adaptability, and genetic programmability of proteins, providing a powerful platform for challenging chemical transformations (*3*).

Despite the development of a variety of metal-binding protein design strategies—where metal ions are directly coordinated by amino acid side chains rather than mediated by organic cofactors—the full catalytic potential of de novo metalloenzymes remains largely unexplored (*3–6*). Most prior efforts have focused on constructing novel protein scaffolds capable of metal binding. For mononuclear metal centers—the focus of this work—prior art has predominantly relied on simplified His-His-His coordination motifs with idealized tetrahedral geometries (*3–5, 7*). As a result, only a limited range of reactions has been demonstrated, and few have yet matched the performance of natural enzymes in the absence of directed evolution.

The idealized His-His-His tetrahedral geometry is largely specific for zinc, and chemistry at these sites has focused on hydrolysis, mirroring the well-studied mechanisms of natural metallohydrolases (*5*). Hartwig and co-workers showed that the His-His-His zinc site of carbonic anhydrase can also reduce ketones to alcohols, and a de novo zinc-binding protein was later reported to catalyze the same reaction, though with low efficiency and selectivity (*8, 9*). Installing a guiding residue during RFDiffusion-based design improves performance substantially, but requires prior mechanistic insight (*10*).

A more challenging and still unrealized goal is the design of highly efficient enzymes built on earth-abundant transition metals other than zinc—iron, nickel, copper, and manganese, which open up a far wider range of chemistry. These metals require not only the near-ideal tetrahedral arrangement seen with zinc, but also octahedral, square-planar, and other coordination geometries (*11*). Using iron as an example, which offers far richer catalytic potential than zinc: natural iron enzymes catalyze hydroxylation of unactivated C–H bonds (α-ketoglutarate-dependent dioxygenases) (*12, 13*), cleave aromatic rings (catechol dioxygenases) (*14*), as well as other reactions that underpin both primary metabolism and natural product biosynthesis (*15, 16*). Natural iron enzymes mostly use octahedral coordination, imposes tighter geometric constraints on the supporting backbone than the tetrahedral sites of zinc enzymes (*3, 6*). New iron catalysis has yet largely relied on repurposing natural enzymes, as exemplified by the iron-catalyzed alkene difunctionalization recently reported by Huang, Jia, and co-workers (*17, 18*). In such cases the parent natural enzyme typically shows low yield and selectivity until extensive directed evolution is applied.

A central challenge in de novo enzyme design is the creation of catalysts for reactions that reach yield and selectivity beyond those obtained by directed evolution of natural enzymes (*19, 20*). This gap is especially wide for metalloenzymes, whose catalytic potential far exceeds what current design methods can access. In this study, we developed a pipeline that produces efficient metalloenzymes directly from computation, targeting two most representative coordination geometries: tetrahedral for zinc and octahedral for iron. We introduced Metal Mer, a structural unit implemented in the software platform METALPROT, for mononuclear metalloenzyme design. We showed that enzymes designed by our approach rival or surpass natural scaffolds in both reactivity and selectivity, for zinc-catalyzed ketone reduction and for iron-catalyzed alkene trifluoromethylazidation.

### The Metal Mer structural unit and METALPROT

The Protein Data Bank provides a vast record of experimentally observed protein–metal interaction geometries, and sampling directly from this distribution is a natural basis for designing metal-binding proteins (*21*). This led us to create the Metal Mer–a structural unit that maps protein backbone to statistically preferred positions of bound metals (Fig. 1a).

**Fig. 1.**
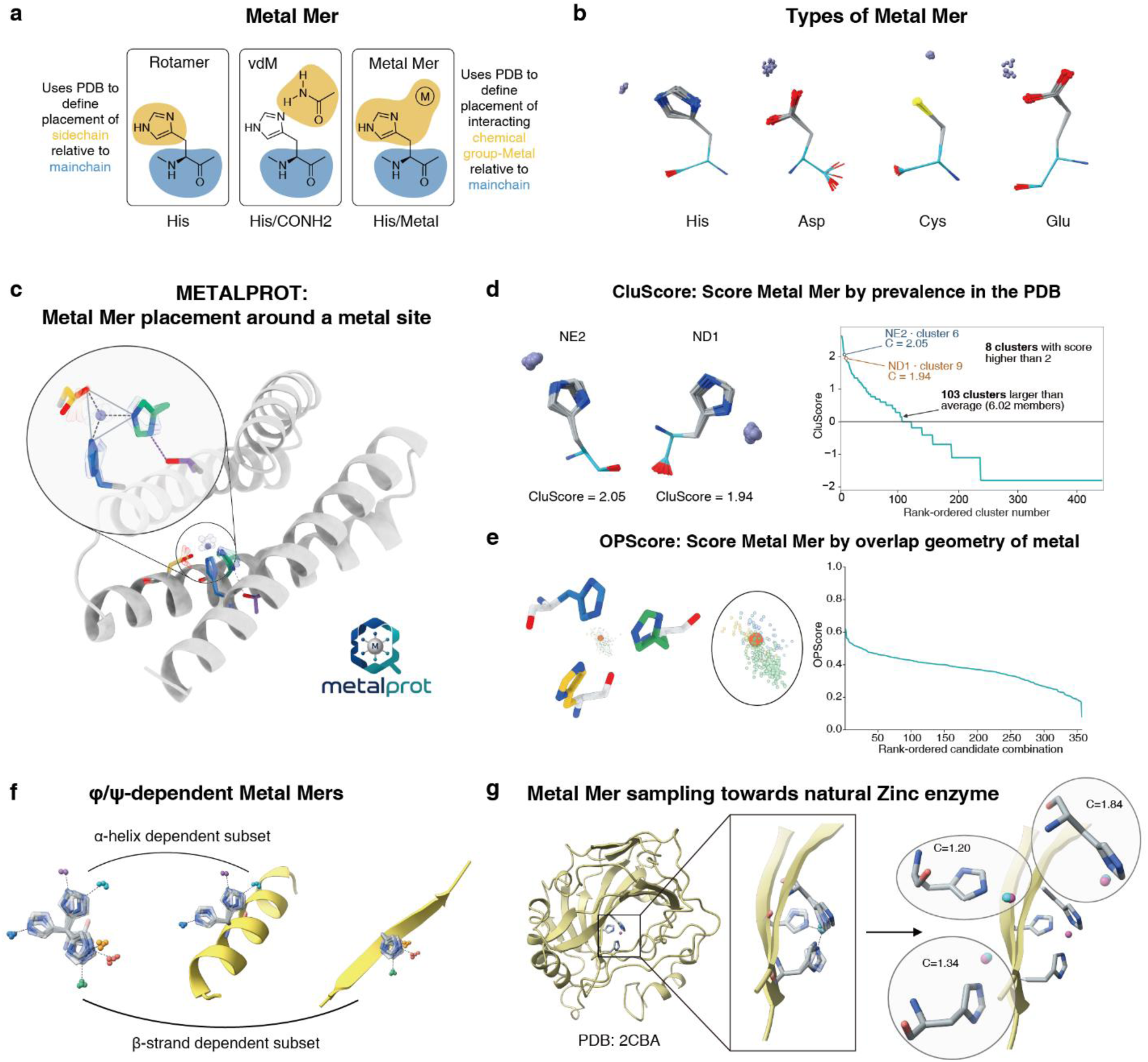
Overview of Metal Mers and METALPROT. (**A**) A Metal Mer is a structural unit that maps protein backbone to statistically preferred positions of bound metals. (**B**) Metal Mers are classified into four types according to their residue-specific preferences. (**C**) METALPROT aligns Metal Mers to a target backbone and then uses a graph-based search to identify combinations of mutually compatible Metal Mers. (**D**) Metal Mers are ranked by prevalence in the PDB, quantified by a cluster score (CluScore); two clusters of His/Zn Metal Mers are shown as an example. (**E**) An overlap score (OPScore) quantifies the spatial convergence of candidate Metal Mers by measuring their collective support for a common metal position. (**F**) Metal Mer clusters show distinct preferences in backbone φ/ψ space. (**G**) Reconstruction of the metal-binding site in carbonic anhydrase II (hCAII; PDB: 2CBA). Metal Mers are sampled on the hCAII backbone to generate possible locations for productive interactions with the chemical groups; here, three clusters of His/Zn Metal Mers are sampled at three positions of the backbone. C denotes CluScore.

Metal Mer extends the van der Mer (vdM) concept of DeGrado and colleagues, which is a statistically weighted unit that record where a protein backbone sits relative to an interacting chemical group (*22*). Using these units to identify potential protein-ligand interactions directly from backbone positions has proven effective in several design settings, including binders for the anticoagulant apixaban and for poly (ADP-ribose) polymerase inhibitors (*22, 23*). We reasoned that a metal-specific implementation of this framework could provide a general route to the design of metal-binding proteins and metalloenzymes.

Starting from constructing a database of metalloprotein structures, we extracted local coordination motifs and perform statistical clustering to generate a library of single-residue metal binding units annotated with cluster-level information (Fig. S1). The resulting Metal Mer library captured residue preferences and metal-dependent distributions of first-shell coordination distances and angles for His, Asp, Glu and Cys (Fig. 1b, Fig. S2). These distributions encompassed characteristic tetrahedral, octahedral and other coordination geometries. These observations provided a structural basis for sampling residue-metal arrangements and defining geometric constraints for subsequent site combinations.

We then ranked Metal Mer clusters using a cluster score (CluScore) derived from their membership counts. This score describes a motif’s relative prevalence within the database and its specified statistical reference (Fig. S3). Two His-Zn representative clusters with CluScore of approximately 2 captured coordination through the imidazole Nε2 and Nδ1 atoms, respectively (Fig. 1d). Combining a binding site requires coordinating side chains from different scaffold positions to converge on a shared metal center. We further introduced an overlap score (OPScore) to quantify the spatial convergence of the Metal Mers generated from different backbone positions (Fig. 1e), where the metal density is calculated in the overlapping region (Fig. S5). Whereas CluScore describes the prevalence of an individual motif cluster, OPScore measures collective spatial support for a shared metal location.

Metal Mer clusters also exhibited distinct backbone φ/ψ preferences, linking their local coordination motifs to the conformation of the target scaffold. Representative motifs from α-helical and β-sheet environments differed in side-chain conformation and metal position relative to the backbone (Fig. 1f). We therefore considered backbone compatibility alongside cluster prevalence when selecting side-chain-metal configurations for individual scaffold positions (Fig. S4).

We implemented this Metal Mer workflow in an end-to-end search framework, METALPROT (Fig. 1c). METALPROT will align the Metal Mers onto a target backbone, and then apply a graph-based search algorithm to identify triplets of pairwise compatible Metal Mers (Fig. S6). Candidate combinations are subsequently ranked according to cluster size, local metal overlap density, and geometric fit, yielding prioritized designs for local metal-binding sites (Fig. 1c). METALPROT samples the positions of coordinating side chains relative to the backbone from experimentally observed structural distributions, thereby enriching for geometries compatible with productive metal binding and potentially with diverse modes of catalysis.

METALPROT can be applied to any protein backbone, provided that there is sufficient space to install the metal-binding residues. To test whether these local motifs recover native coordination geometries, we reconstructed the zinc site of human carbonic anhydrase II (hCAII, PDB: 2CBA). The reconstructed metal positions and coordinating side-chain arrangements resembled those in the native structure, which supports the use of Metal Mers for assembling local coordination environments on a specified backbone (Fig. 1g, Fig. S7).

### Computational workflow of de novo metalloenzyme design

We next assembled METALPROT into a complete design pipeline, from backbone generation to sequence design. We observed that the recent studies from Baker and colleagues, which examined a wide range of backbones, found that simple helical bundles far out-performed more complex architectures in Zn-binding esterase design and the most active architecture was a 5-helix bundle (*5*). Others showed impressive progress with de novo 4-helix zinc enzymes or bimetallic proteins (*6, 24, 25*). Moreover, the stability of helical bundles is impressive for new-to-nature reaction catalysis (*20*). Thus, we focused our efforts on generating 4- and 5-helix bundles, although the method can be generalized to any backbone.

We first assembled a set of 4-helix and 5-helix bundles, which were drawn from two sources:

(i) previously reported backbones known to bind small molecules since they already support stable folding, and (ii) parametrically generated helical bundles designed to sample a broader geometric space (Materials and methods)(*26*). To maximize structural diversity, we introduced additional parameters controlling helix placement and tilt, allowing bundles with funnel-shaped, rectangular, and other non-canonical cross-sections to be generated (Fig. 2a, S8). After iterative sequence design and clustering, we obtained a robust ensemble of helix-bundle backbones that served as a template library for Metal Mer placement.

**Fig. 2.**
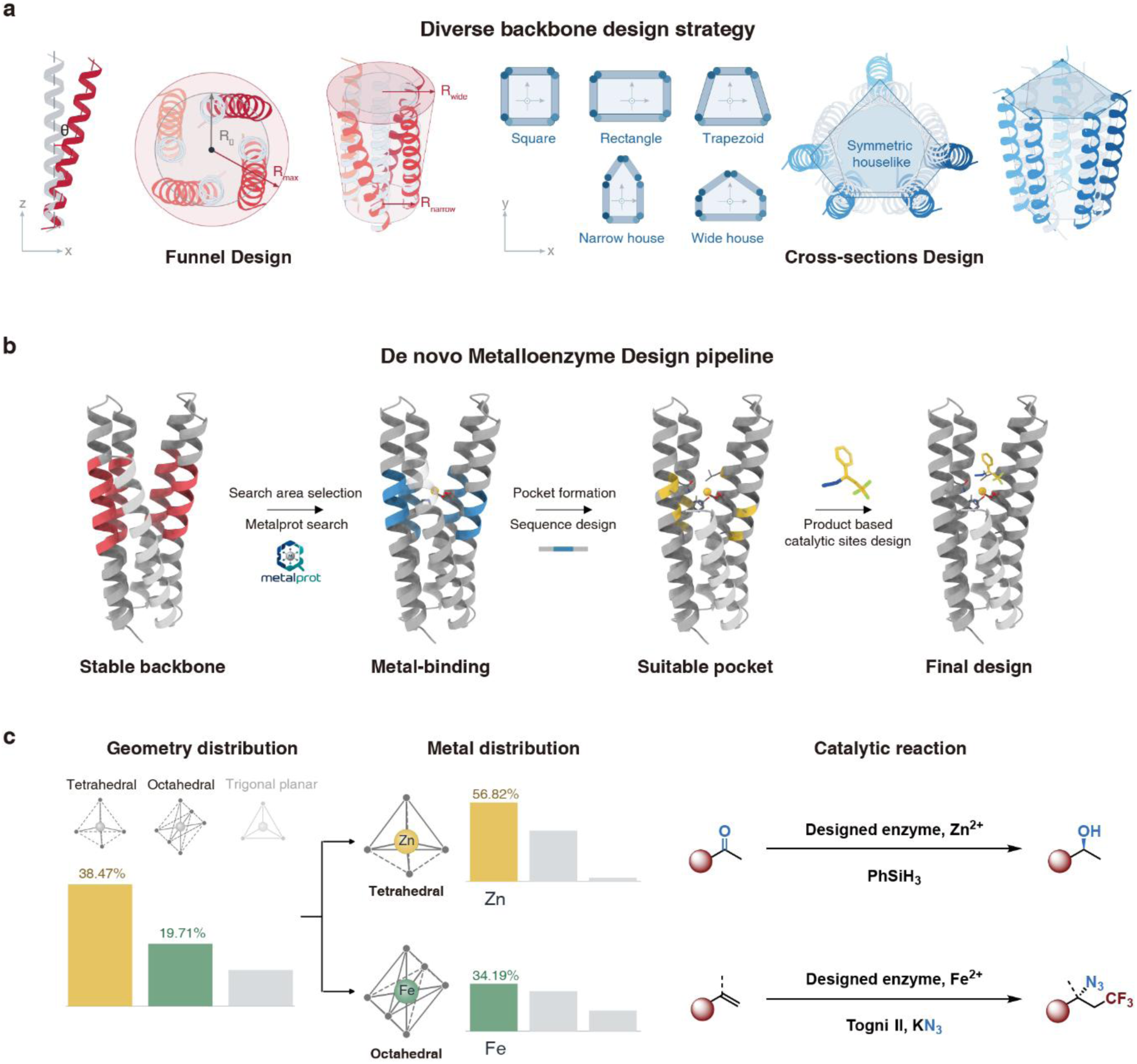
Metalloenzyme Design Strategy. (**A**) Diversifying backbones through funnel design and cross-section design. (a) Left, schematic of helix tilting in the x-z plane. Middle, superposition of the reference (gray) and modified(red) backbones viewed along z. Right, oblique view with an illustrative geometry. θ denotes the helix-axis tilt; R_0_ and R_max_ indicate the projected radii of the reference and modified bundles, respectively; R_narrow_ and R_wide_ denote the narrow-end radius, wide-end radius and axial length of the illustrative geometry. (b) Left, schematic square, rectangular, trapezoidal, narrow-house and wide-house geometries. Middle, comparison of a regular pentagonal reference arrangement (gray) with a houselike arrangement (blue), viewed along z. Right, an oblique view highlights changes in helix arrangement and cross-sectional geometry. (**B**) De novo metalloenzyme design pipeline. Red, blue and yellow regions indicate sampling windows, local sequence-design regions and selected sidechain positions, respectively. (**C**) Coordination geometry and metal distributions with corresponding catalytic reaction designs. The analysis comprises 7,507 coordination geometry counts from MetalPDB(*45, 46*). Tetrahedral and octahedral geometries rank first and second in abundance, respectively, accounting for 2,888/7,507 (38.47%) and 1,480/7,507 (19.71%) of the total. Within these geometry categories, Zn is the most frequently represented metal in tetrahedral coordination (1,641/2,888; 56.82%), while Fe is the most frequently represented metal in octahedral coordination (506/1,480; 34.19%). The reaction schemes on the right depict catalytic transformations designed for the corresponding Zn- and Fe-containing enzymes.

We next used METALPROT to place Metal Mers and corresponding metal sites onto these backbone templates. This search produced multiple combinations of coordinating residues (Fig. 2b). We retained only those Metal Mers positioned within the bundle interior and carved out a rudimentary catalytic pocket via adjustment of the scaffold around the predicted metal site to facilitate substrate ingress and product egress (Materials and methods).

These initial metal-binding proteins then underwent multiple rounds of iterative design, each with a distinct objective: (i) establishing a high-confidence, designable, and geometrically stable first-shell coordination environment; (ii) introducing at least one supporting second-shell interaction; and (iii) passing a series of computational filters followed by fine adjustment of pocket volume (Fig. 2b). In the final step, a single product enantiomer is used as the selectivity template, placed into the catalytic cavity with transition state (TS)-like distance and angle restraints (Fig. 2b). We carried out constrained sequence design using Rosetta with side-chain repacking and minimization to organize favorable interactions around the restrained product (Materials and methods). This shifts the focus from TS search to candidate validation. Because only one product configuration is used to generate the pocket, the resulting design is biased toward that enantiomer if the enantio-determining step directly leads to the product. Hammond’s postulate suggests that product-like structure can be a useful initial approximation for TS (*27*). Because the product lacks geometric features from TS, established restraints on bond formation/cleavage, attack angle, etc. are used to introduce them. This restrained-product strategy is a design hypothesis for filtering geometrically and electronically plausible enantioselective enzyme candidates (Fig. S10b).

We applied this approach to two catalytic systems representing the two most common coordination geometries observed in nature: a zinc site with tetrahedral coordination and an iron site with octahedral coordination, each designed for a different reaction (Fig. 2c). Both systems use the same combinatorial search framework, with geometric constraints tailored to their respective coordination requirements (Materials and methods).

### Artificial Zn Enzyme for Enantioselective Hydride Transfer

The zinc enzyme Human Carbonic Anhydrase II (hCAII) can catalyze a stepwise hydride-transfer reaction: a silane such as PhSiH₃ first reacts with the zinc hydroxide to generate a highly reactive enzymatic zinc hydride, which delivers H^−^ to the carbonyl with stereochemical control from the pocket environment (*8*). In hCAII, the zinc is coordinated by three histidines in near-ideal tetrahedral geometry. T199 of hCAII additionally serves both as an electrophilicity promoter and as a substrate-binding anchor, and thus been incorporated into de novo designs to achieve high activity (*8, 10*). In our designs, we deliberately sampled alternative coordinating residues and omitted the hydrogen-bonding residue (Fig. 3a). Our aim was to test whether a minimal, stochastically sampled combination of coordinating residues—without prior knowledge of the native catalytic pocket—could provide accurate design and support catalysis.

**Fig. 3.**
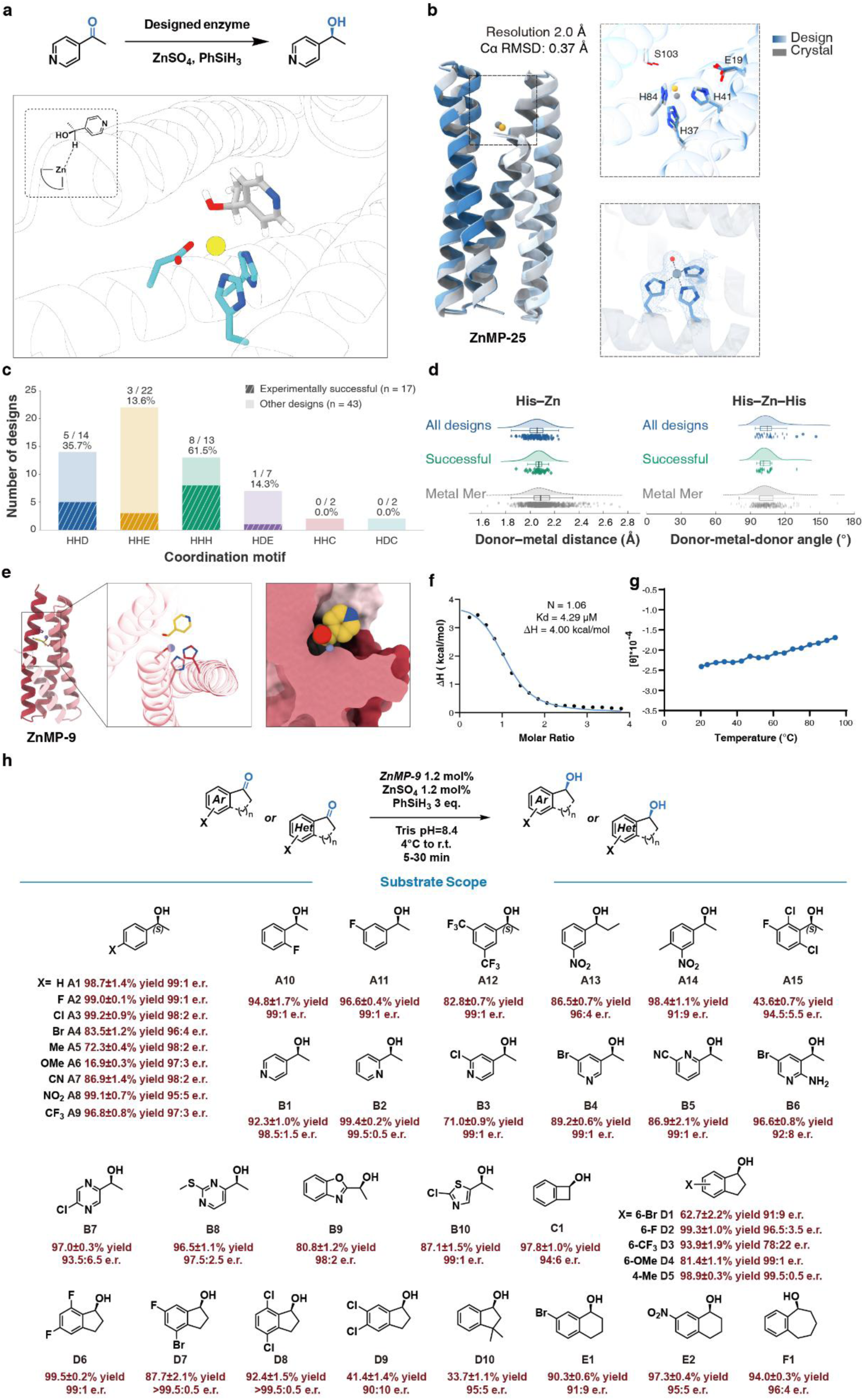
Zinc Hydride Transferase. (**A**) Applying restrained product strategy to model substrate and the resulting structure. (**B**) Single-crystal X-ray diffraction results. (Upper) Aligned ZnMP-25 crystal structure (PDB code 47PF) and AF3 predicted model. (Lower) Zinc-binding site of ZnMP-25. The 2Fo–Fc electron density map (blue mesh) is contoured at 1.0 σ. The zinc ion is shown as a blue sphere, water shown in red, coordinating residues are shown as sticks. (**C**) Analysis of experimentally successful designs grouped by coordination motif. Stacked bar chart showing the total number of designs for each motif (HHD, HHE, HHH, HDE, HHC, HDC). Bars are subdivided into experimentally successful designs (hatched, n = 17) and other designs (solid, n = 43). The number of successful designs out of the total tested for each motif, along with the success percentage, is displayed above each bar. (**D**) Distributions of His–Zn coordination distances (left) and His–Zn–His coordination angles (right). Distances are measured between His donor nitrogen atoms and Zn; angles are defined by donor nitrogen atoms from two distinct His residues, with Zn at the vertex. Blue, teal and grey indicate all designs, the experimentally successful subset and Zn Metal Mer reference cores, respectively. His ND1/NE2 coordination is pooled across coordination motifs. Curves show kernel density estimates, and points represent individual distance or angle observations. (**E**) Substrate binding pose in ZnMP-9. (**F**) Isothermal titration calorimetry (ITC) analysis of ZnSO_4_ binding to ZnMP-9. ITC experiments were performed in 50 mM MES pH 6.0, 150mM NaCl at 25 °C. Integrated heat data plotted as a function of the molar ratio of ZnSO_4_ to ZnMP-9. The blue line represents the best fit to a one-site binding model, yielding N = 1.06, K_d_ = 4.29 μM, and ΔH = 4.00 kcal/mol. (**G**) Circular dichroism spectra of holo-ZnMP-9 at varying temperatures. (**H**) Substrate scope of ZnMP-9 (2 µmol scale). Yields were determined with GC-MS under single-ion monitoring mode using 1,3,5-trimethoxybenzene as internal standard. Enantiomeric excess was determined by chiral HPLC or chiral GC. All experiments were carried out in triplicates.

We initially designed 60 zinc enzymes using the Metal Mer strategy with 4-acetylpyridine as the model substrate, 20 were built on four-helix bundles and 40 on five-helix bundles (Table S3). To test the accuracy of our design method, we crystallized ZnMP-25, a five-helix bundle design, with 2.0 Å resolution (Fig. 3b). The structure shows that the pipeline produces zinc-binding proteins closely matching the design model, with a global backbone Cα RMSD of

0.37 Å. Clear electron density for the bound zinc ion, a coordinating water, and the three binding residues together confirmed the tetrahedral metal coordination geometry highly consistent to the designed model. Moreover, the second-shell ligands (Glu19, Ser103) are as designed (Fig. 3b). Such precision will be needed in applications where a specific ligand geometry is used to influence reactivity. This compares favorably with a diffusion-based workflow showing discrepancy between crystal structure and designed model on one of the zinc-coordinating histidines (*5*). Thus, the Metal Mer strategy provides highly precise designs when compared to other state-of-the-art methods.

We tested the designs against the model substrate 4-acetylpyridine, assaying formation of the corresponding alcohol by GC-MS (Table S4). Seventeen of the 60 designs showed catalytic activity, and three delivered above 95:5 e.r.. Active designs span both 4- and 5-helix bundles, confirming that Metal Mer sites are not confined to a single fold. Activity also arose across several coordination modes, including His-His-Asp and His-His-Glu (Fig. 3c). Moreover, the designed zinc sites are not confined to ideal tetrahedral geometry: in the design models, the residue-to-zinc distances and the residue–zinc–residue angles span the range observed across natural metalloproteins (Fig. 3d).

Among the 60 designs screened, variant ZnMP-9 consistently outperformed the others. In the design model an S-configuration product was picked as a post-transition-state geometry by restraining the ZnH distance. ZnMP-9 coordinates zinc through an Asp-His-His triad, notably different from the three-histidine site of hCAII and of previously reported de novo designs (Fig. 3e). In the absence of hydrogen-bond activating residues, ZnMP-9 and several other active variants shows that, such residues conserved in hCAII and preserved in previous design work is not required for efficient catalysis.

CD spectra indicate that ZnMP-9 is α-helical, consistent with the designed bundle (Fig. S25). The protein is also highly thermostable: helicity is largely retained on heating to 90°C (Fig. 3f). Zn(II) binding was confirmed by native mass spectrometry, and ITC titrations of Zn(II) into ZnMP-9, ZnMP′-6, and ZnMP′-9 confirmed 1:1 binding with K_d_ values of 4.29, 0.985, and 0.676 µM, respectively (Fig. 3f, S27b).

The ZnMP-9/ZnSO₄/PhSiH₃ system reduces a diverse scope of aryl- and heteroaryl- ketones under mild conditions (Fig. 3g). Para-substituted acetophenones are converted with up to 99.1% yield and high enantioselectivities (95:5 to 99:1 e.r.); electron-donating Me and OMe groups lower yields while preserving high stereocontrol (98:2 and 97:3 e.r.). The scope extends to ortho- and meta-substituted aryl ketones (A10-A11), an aprepitant fragment (A12, 82.8% yield, 99:1 e.r.), and a crizotinib fragment (A15, 43.6% yield, 95:5 e.r.), demonstrating tolerance of drug-like functionality. Heteroaryl methyl ketones (B1-B10), including pyridine, pyrimidine, pyrazine, benzoxazole, and thiazole derivatives, are reduced with consistently high enantioselectivity (up to 99.5:0.5 e.r.) and yields up to 99.4%, despite the presence of coordinating heteroatoms and diverse ring electronics. Fused bicyclic ketones (C1, D1–D10, E1–E2, F1) extend the scope to benzocyclobutenols and substituted indanols, with several examples exceeding 99.5:0.5 e.r. (D4–D8) and yields up to 99.5%; more challenging cases (D3, 78:22 e.r.; D9, 90:10 e.r.; D10, 95:5 e.r.) define the current limits.

### Functionalization of Complex Natural Products and Drug-Like Scaffolds

Chiral alcohols are common among FDA-approved drugs, yet their synthesis via asymmetric reduction remains challenging (*28, 29*), particularly when a complex scaffold contains multiple carbonyl groups of similar reactivity. De novo metalloenzymes have not yet demonstrated selective catalysis on complex, polyfunctional substrates: their limited pocket size constrains reactivity toward large substrates, and selectivity decreases with heavily substituted substrates. We envision that the binding pocket of a de novo helical bundle can be reshaped independently of the catalytic residues, adapting it to larger, more demanding substrates (Fig. 4a).

**Fig. 4.**
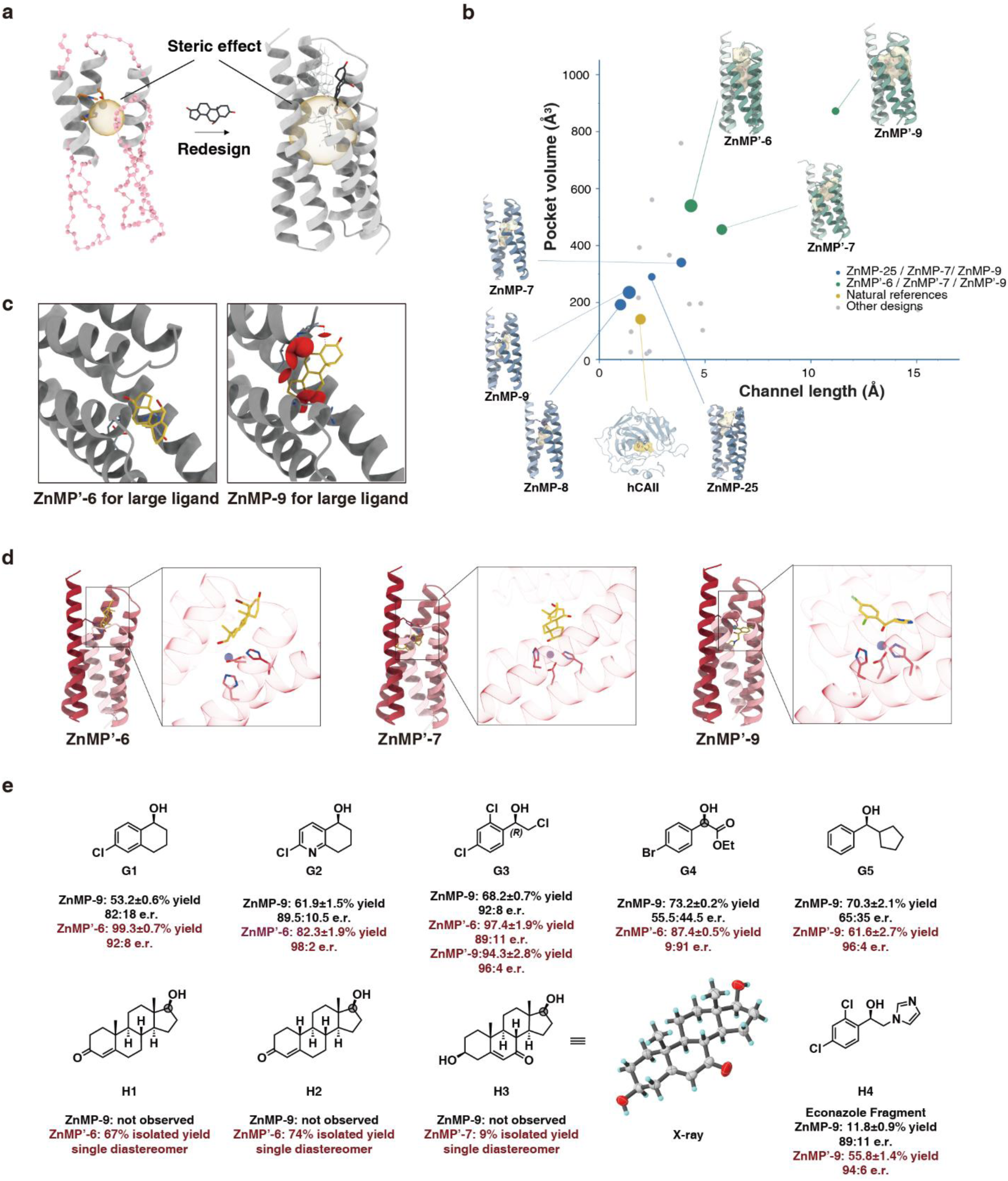
Redesign of Zinc Hydride Transferase for Challenging Substrates. (**A**) Partial diffusion generates active enzymes with enlarged pockets. (**B**) Analysis of channel length and pocket volume for designed zinc metalloproteins compared to natural references. Scatter plot comparing the channel length (Å) on the x-axis against the pocket volume (Å³) on the y-axis. Data points were calculated using CAVER. Points are color-coded according to the legend: blue represents ZnMP-25, ZnMP-7, and ZnMP-9; green represents ZnMP′-6, ZnMP′-7, and ZnMP′-9; yellow indicates the natural reference hCAII; and grey represents other designs or natural references. (**C**) A triterpene substrate binding pose with same restraints lead to higher steric clashes in ZnMP-9 than expanded ZnMP′-6. (**D**) Substrate binding pose in ZnMP′-6, ZnMP′-7, and ZnMP′-9. (**E**) Scope of challenging substrates. Site selectivity indicated by grey circle. Yields (2 µmol scale) were determined with GC-MS under single-ion monitoring mode using 1,3,5-trimethoxybenzene as internal standard. Enantiomeric excess was determined by chiral HPLC or chiral GC. Experiments were carried out in triplicates. For H1-H3, isolated yield was determined on 0.1 mmol scale.

Applying to ZnMP-9, we kept the zinc binding sites and short helixes supporting each residue, using partial diffusion in RFDiffusion (*30*) to generate longer designs (Materials and methods). After filtering the shapes, we obtained a series of new helix bundles with an enlarged pocket and additional residues available for substrate contact. Sequences were then refined using the desired product isomer with our restrained product analog strategy, with selectivity embedded in the design.

With this procedure, we tested 11 of the improved de novo designs, all of which were active, as compared to 17/60 from the initial design. The zinc-binding residues are structurally unchanged, whereas the active-site pockets differ substantially in volume and depth: the first-round designs are comparable to hCAII, while the redesigns are roughly 2- to 4-fold larger (measured with CAVER (*31*)) (Fig. 4b), with intrinsic shape diversity.

Three of these, ZnMP′-6, ZnMP′-7 and ZnMP′-9, convert bulkier substrates with substantially improved stereoselectivity (Fig. 4d). The G and H series include densely substituted ketones such as steroidal scaffolds and drug-fragment ketones (Fig. 4e).

Additional rings and functional groups not only increase 3-dimensional steric repulsion but also introduce competing coordination or alternative binding poses that erode enantioselectivity. To our delight, for substrates with increased sterics around para- and alpha-position to carbonyl group, such as G1–G3, the redesigned variants improve both yield and enantioselectivity. For G4, ZnMP-9 gives only 56:44 e.r., whereas ZnMP′-6 gives 9:91 e.r. with reversed chirality, likely due to ester-specific interactions. For G5 with comparable size on either side of the carbonyl, differentiation between phenyl and cyclopentyl significantly improves from 65:35 e.r. (ZnMP-9) to 96:4 e.r. (ZnMP′-6).

We next tested the designs on the conversion of androstenedione to testosterone, a key transformation in steroid metabolism and industrial production (*32, 33*). The synthesis requires selective reduction of the C17 ketone to a 17β-hydroxyl while leaving the conjugated C3 ketone intact, which is essential to the pharmacophore (*34*). In our design, ZnMP′-6 presents only the C17 ketone to the metal hydride, delivering testosterone (H1) with >19:1 site- and diastereoselectivity in 67% isolated yield without protecting groups. H2 was similarly converted (74% isolated yield, single diastereomer). For H3 bearing drastically different substitution pattern, variant ZnMP′-7 selectively afforded the 21β-hydroxyl product as a single diastereomer, confirmed by its crystal structure. The parent ZnMP-9 gives no detectable conversion of all tested steroid substrates.

### Design of Artificial Iron Enzyme for Alkene Difunctionalization

Iron’s Fe(II)/Fe(III) redox cycle supports one-electron catalysis, but the radical intermediates it generates are short-lived, making selectivity control a central challenge (*35, 36*). We targeted alkene trifluoromethylazidation to install two valuable pharmacophores in one step: CF₃ enhances lipophilicity and metabolic stability, while N₃ provides a handle for click chemistry (*37, 38*). This reaction involves three components—an alkene, a CF₃ radical source, and an azide—and orchestrating their sequential transformation while controlling the fate of transient radical intermediates within a single enzyme active site posts a significant challenge to enzymatic catalysis. Two non-heme iron enzymes, AoHMS and BsQueD, have recently been engineered to catalyze this reaction, but only after more than ten rounds of directed evolution to reach 73% and 70% yields, 96:4 and 98:2 e.r., respectively (*17, 18*).

The proposed reaction mechanism starts a one-electron reduction of hypervalent iodine reagent by the Fe(II) center, generating a trifluoromethyl radical that undergoes addition to the alkene double bond. The resulting alkyl radical can further react with an iron-bound azide in a radical-rebound fashion to form the chiral alkyl azide product (Fig. 5a) (*39, 40*).

**Fig. 5.**
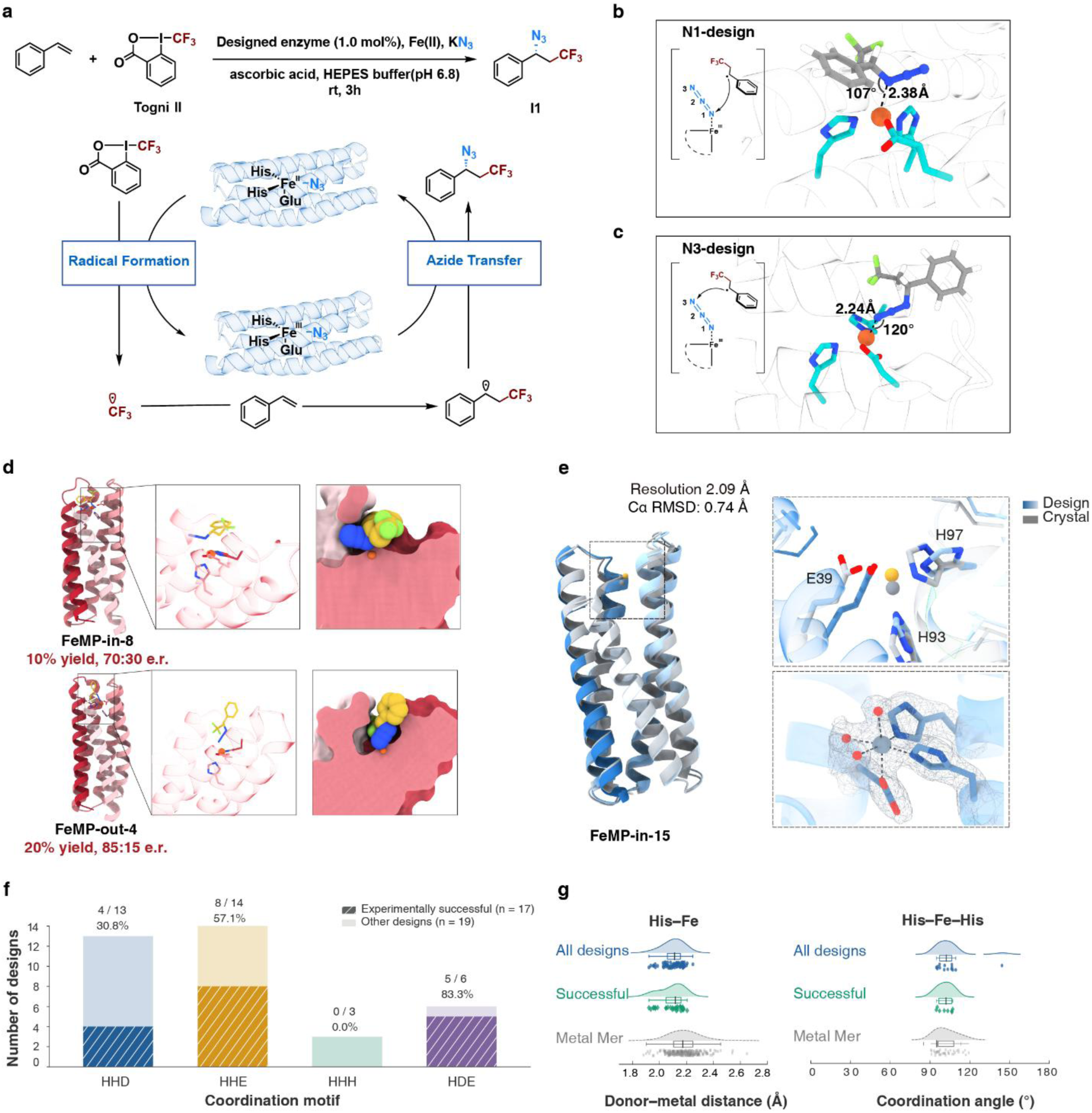
Nonheme Iron Enzyme Design for Alkene Trifluoromethylazidation. (**A**) Model reaction Scheme. Standard conditions: Fe(II) (1 mM), NaN_3_ (50 mM), substrate (2.5 mM), Togni’s reagent II (5 mM), ascorbic acid (0.1 mol%), HEPES buffer (50 mM, pH 6.8), and purified enzyme (1.0 mol%). (**B**) N1-selective C-N bond formation geometry and design parameters. (**C**) N3-selective C-N bond formation geometry and design parameters. (**D**) FeMP-in-8 and FeMP-out-4: design and experimental results from reaction with styrene. (**E**) Single-crystal X-ray diffraction results. (Upper) Aligned FeMP-in-15 crystal structure (PDB code 47HF) and AF3 predicted model. (Lower) Metal binding site of FeMP-in-15. The 2Fo–Fc electron density map (grey mesh) is contoured at 1.0 σ. The cobalt ion is shown as a blue sphere, water shown in red, coordinating residues are shown as sticks. (**F**) Analysis of experimentally successful iron enzymes grouped by coordination motif. Stacked bar chart showing the total number of designs for each motif (HHD, HHE, HHH, HDE). Bars are subdivided into experimentally successful designs (hatched, n = 17) and other designs (solid, n = 19). The number of successful designs out of the total tested for each motif, along with the success percentage, is displayed above each bar. (**G**) Distributions of His–Fe coordination distances (left) and His–Fe–His coordination angles (right). Distances are measured between His donor nitrogen atoms and Fe; angles are defined by donor nitrogen atoms from two distinct His residues, with Fe at the vertex. Blue, teal and grey indicate all designs, the experimentally successful subset and Fe Metal Mer reference cores, respectively. His ND1/NE2 coordination types are combined, and observations are pooled across coordination motifs. Curves show kernel density estimates, and points represent individual distance or angle observations.

We extended the Metal Mer approach to construct iron sites for this reaction. The iron site was constrained to octahedral geometry, and the first-shell ligand composition was left unbiased, as for the zinc designs. Rather than modeling a transition state containing all three reaction components, our product-based strategy used (1-azido-3,3,3-trifluoropropyl)benzene, the reaction product, as the target for sequence refinement.

Following previous studies, we assumed a radical pathway in our design: once the trifluoromethyl radical adds to the alkene, coupling with iron-bound azide can occur at either N1 or N3 position, plausibly influencing enantioselectivity. Therefore, N1 and N3 attack were treated as alternative design hypotheses rather than as a settled mechanism (Fig. 5b, 5c).

We initially reasoned that the iron-coordinated N1 would be more activated toward radical attack, so we took previously reported computational parameters for analogous C–N_3_ bond formation as reference values, and employed Rosetta (*30*) to run N1 designs (*41, 42*). In the N1 design, the Fe-N1 distance and Fe-N1-C1 angle were constrained to 2.38 Å and 107° (Fig. 5b). We initially built 30 designs around an N1-attack mechanism, in which the benzylic radical bonds directly to the azide nitrogen coordinated to iron. Of the 30 designs, 12 gave product detectable by GC-MS, in which FeMP-in-8 and FeMP-in-15 have a moderate 10% yield and the e.r. achieved 70:30 (Table S7).

Given the modest enantioselectivity of the N1-attack designs, we reasoned that these geometric parameters may be insufficient to enforce effective stereocontrol. To circumvent this, we used the same computational workflow to design 6 additional variants based on an N3-attack geometry. In the N3 design, the Fe-N1 distance and Fe-N1-N2 angle were constrained to 2.24 Å and 120° (Fig. 5c). Five N3-attack designs were active, and the best, FeMP-out-4, gave 85:15 e.r.—exceeding the best N1-attack designs (Fig. 5d). These results do not establish which pathway operates in the enzymes, but they indicate that the N3-attack geometry is more readily accommodated in a designed pocket. Consistent with this, redesigning the backbone of FeMP-in-15 did not improve enantioselectivity, suggesting that the limitation lies in the N1-attack geometry itself rather than in the surrounding environment (Table S6, S7).

To confirm that the octahedral iron geometry was realized as designed, we screened several designs for crystallization and obtained a structure of FeMP-in-15, with cobalt substituted for iron—a common surrogate that adopts comparable coordination geometry and is more stable under crystallization conditions (*43, 44*). The structure agrees closely with the AF3-predicted design model, with a backbone Cα RMSD of 0.74 Å (Fig. 5e). The coordination geometry is precisely octahedral as we designed, with well-defined electron density for the bound metal ion, three coordinating waters, and the three monodentate protein ligands with the correct ligand-metal distances. The side chain of the iron coordinating glutamate shifts slightly relative to the design model, possibly owing to the Co substitution. Overall, this showed that the selected Metal Mers are highly precise not only for zinc enzyme design but could also be extended to the precise design of other metalloenzymes.

Across all 30 N1-attack and 6 N3-attack designs, 17 were active directly from computation (Table S7). In the success designs, we observed diverse first-shell compositions, as for the zinc enzymes. The natural enzymes reported to catalyze this reaction use His-His-Glu and His-His-His-Glu motifs, whereas our active designs span His-His-Glu, His-His-Asp, and His-Asp-Glu motifs (Fig. 5f). The octahedral geometries extended beyond those of the two reported natural enzymes, spanning the range of distances and angles observed across natural metalloproteins as a whole (Fig. 5g).

Our best design, FeMP-out-4, approaches but does not yet match the performance of directed-evolution-optimized natural enzymes. To identify the bottleneck for further improvement, we examined its design model. The pocket structure suggested that Ala116 leaves sufficient room for the intermediate to approach N3 in more than one conformation (Fig. 6a). We reasoned that a bulkier side chain at this position would restrict access to a single productive geometry. Consistent with this, the single mutation A116I and A116V increased enantioselectivity from 85:15 e.r. to 90:10 e.r. and 93:7 e.r., respectively, approaching the performance of best engineered natural enzymes from directed-evolution (95:5 e.r. and 93:7 e.r.). As expected, introducing a less sterically demanding side chain (A116G) opened additional space in the pocket and lowered the e.r. to 80:20. This structure-activity relationship indicates that, although our pocket design strategy using Rosetta can refine the catalytic sites, it does not yet fully account for the effects of subtle steric and conformational factors that dictate precise enantiocontrol.

**Fig. 6.**
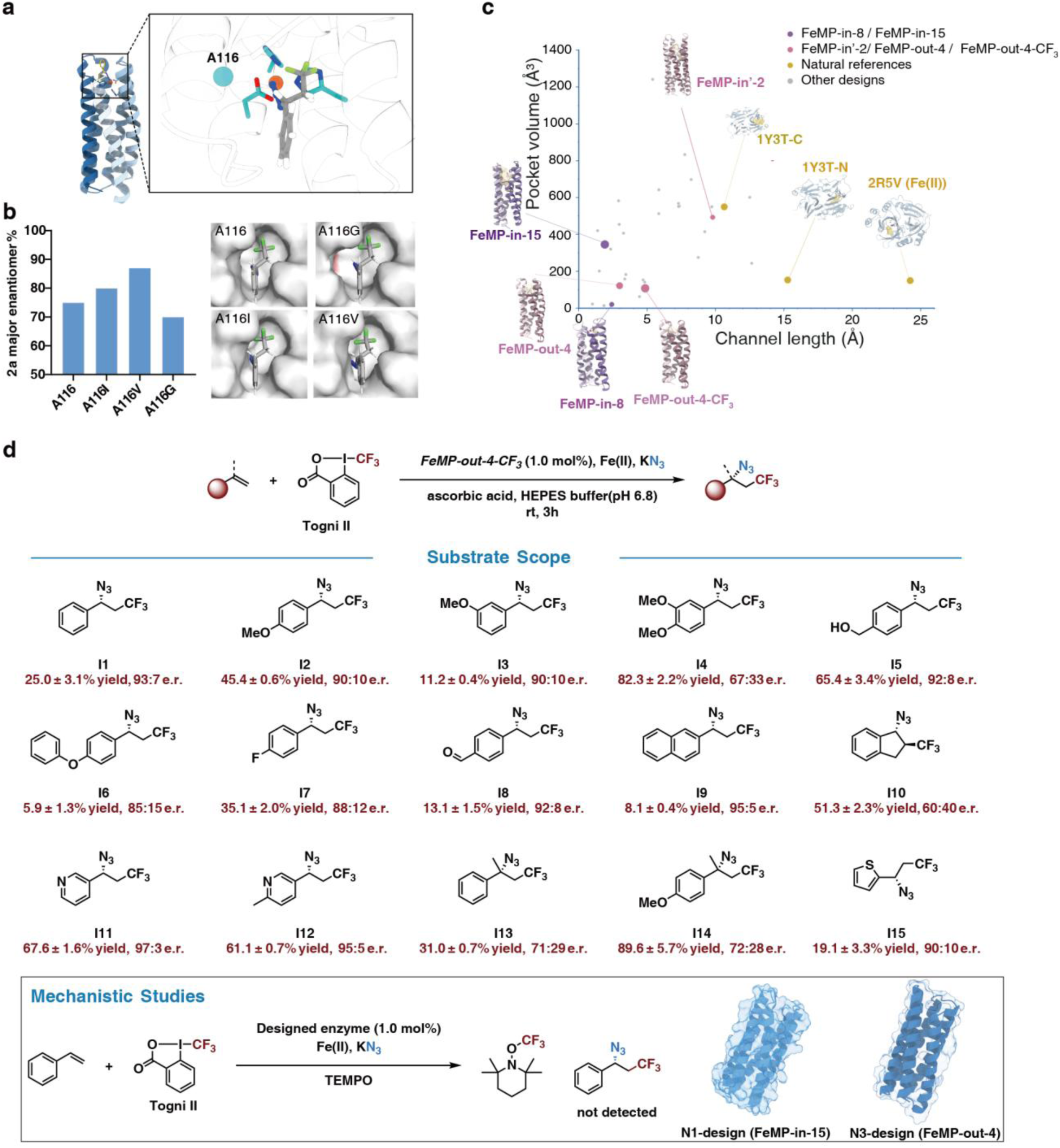
Site-Directed Mutagenesis and Substrate Scope. (**A**) Protein structure of FeMP-out-4. The cyan spheres represent mutated residue-A116. (**B**) Site-directed mutagenesis modulates enantioselectivity and protein cavity architecture. (**C**) Analysis of channel length and pocket volume for designed iron metalloproteins compared to natural references. Scatter plot comparing the channel length (Å) on the x-axis against the pocket volume (Å³) on the y-axis. Data points were calculated using CAVER. (**D**) Substrate scope and mechanistic studies. Yields were determined by GC-MS. Radical-trapping experiments were performed in both N1 and N3 designs.

FeMP-in-8, FeMP-in-15, and FeMP-out-4 comprise only about 150 residues and share a shallower pocket than the natural enzymes (volume 121.8 Å³, channel length 3.0 Å for FeMP-out-4) (Fig. 6c). This contrasts with our initial expectation that a deeper pocket would be necessary to confine the radical intermediate and ensure selectivity.

With this variant FeMP-out-4-CF_3_ (A116V), we next examined the substrate scope of this biocatalytic platform (Fig. 6d). Electron-rich styrenes bearing *para*-methoxy (I2), *meta*-methoxy (I3), and 3,4-dimethoxy (I4) substituents were effectively converted. Chemically sensitive benzylic alcohol was also tolerated (I5, 65% yield, 92:8 e.r.). Electron-poor styrenes, such as *para*-fluoro (I7), and *para*-formyl (I8) substituents were also compatible. Other aryl-alkene substrates, such as naphthalene (I9) and indane (I10), were also effective. Heteroaryl olefins perform particularly well: 2-vinylpyridine (I11, 68% yield, 97:3 e.r.) and 2-methyl-5-vinylpyridine (I12, 61% yield, 95:5 e.r.). α-Methyl styrenes were also converted (I13 and I14). To probe the mechanism of this abiotic transformation, we further performed a radical-trapping experiment using TEMPO in both N1- and N3-design enzymes (Fig. 6d).

The addition of TEMPO completely inhibited product formation, supporting the involvement of radical intermediates and validating our design hypothesis. The successful design of this three-component reaction on an octahedral iron center thus opens access to previously underexplored classes of de novo metalloenzymes.

## Conclusion

Nature provides a rich repertoire of metalloenzymes, particularly those in which metal ions are directly coordinated by amino acid side chains, exhibiting diverse binding geometries and catalytic functions. We developed the Metal Mer framework to address the current limitations of metalloenzyme design, improving success rate, design accuracy, substrate scope, and the range of accessible reactions.

Ultimately, the Metal Mer pipeline delivered high enantioselectivity across diverse prochiral ketones and precise site- and diastereoselectivity on complex natural products in zinc- catalyzed hydride transfer, while successfully orchestrating the three-component complexity of the iron-catalyzed radical reaction. Both the zinc and iron designs use ligand sets and coordination geometries that depart from those of their natural counterparts, showing that de novo scaffolds can reach evolved-enzyme performance without reproducing natural active- site architecture. Along this study, we have used diverse four- and five-helix bundles to build these catalysts, and provide a parametric route to their generation. Helical bundles will not address every design problem, but their utility is considerable: they have served as the principal scaffold for much of the recent progress in de novo metalloenzymes. We anticipate that the approach described here offers a tractable means of constructing such scaffolds, and that it can be extended to other tertiary architectures.

In summary, METALPROT and Metal Mers provide a general platform for the design of mononuclear metalloenzymes. Beyond zinc and iron-dependent reactions demonstrated here, this strategy opens opportunities for designing catalysts incorporating cobalt, nickel, copper, and other metal ions for a broad range of synthetically useful transformations. Moreover, given that nature also employs bimetallic active sites extensively, we anticipate that Metal Mers can be extended to construct such architectures for the exploration of more complex catalytic chemistries. Finally, although currently implemented as an external design framework, Metal Mers are readily transferable to emerging backbone and sequence design paradigms, including diffusion-based methods, thereby providing a modular component for future computational enzyme design platforms.

## Acknowledgments

We gratefully acknowledge the instrumental facilities and technical support provided by Center of Pharmaceutical Technology, School of Pharmaceutical Sciences, Tsinghua University. We thank Technology Center for Protein Sciences, Tsinghua University for technical assistance in X-ray Crystallography. We thank the ID02-Microfocusing X-Ray Protein Crystallography Beamline of High Energy Photon Source (https://cstr.cn/31138.02.HEPS.ID02) and Shanghai Synchrotron Radiation Facility of BL02U1 Macromolecular X-ray Crystallography Beamline (https://cstr.cn/31124.02.SSRF.BL02U1) for providing technical support and assistance in XRD data collection. We gratefully thank Dr. William F. DeGrado, Dr. Nicholas F. Polizzi, Dr. Kaipeng Hou, Dr. Gang Lu for helpful discussion. Lei Lu is supported by a Bayer THU Investigator Awards.

## Funding

National Natural Science Foundation of China Excellent Young Scientists Fund Program 22577059

Natural Science Foundation of Beijing L252189

Tsinghua University, “Dushi Plan”

Tsinghua University, Youth Talent Cultivation Fund

Tsinghua-Peking Center for Life Sciences, Beijing Frontier Research Center for Biological Structure, Cultivation Fund

## Author contributions

Conceptualization: LL

METALPROT software and algorithm: LL, ZR, (chemistry support from CS, JY)

Analytical method and data analysis (GC-MS, chiral GC/HPLC, NMR): CS, JY

Chemical synthesis and enzymatic reactions: CS, JY

Biochemical assays: JY, YG, ZR ITC and CD: CS, YG

Native MS: XZ, YG, LL

X-ray crystallography: CS (experimental support from YG, XZ, JY, ZR)

Visualization: ZR, JY, CS

Funding acquisition: LL

Writing – original draft: LL, CS, ZR, JY

Writing – review & editing: LL, CS, ZR, JY, XZ

## Competing interests

Authors declare that they have no competing interests.

## Data, code, and materials availability

The METALPROT software, including all associated modules, scripts, and documentation, is available at https://github.com/Mituan02/Metalprot. Coordinates and X-ray structure factors for ZnMP-25 and FeMP-in-15 have been deposited at the RCSB Protein Data Bank (PDB) under accession number 47PF and 47HF. Single-crystal X-ray diffraction data for compound H3 have been deposited in the Cambridge Crystallographic Data Centre under accession number CCDC 2595681. All data supporting the findings of this study (SDS-PAGE, SEC, ITC, CD, native-MS, NMR, chiral LC/GC, and GC-MS) are available within the paper and its Supplementary Information.

## Notes

### Competing Interest Statement

The authors have declared no competing interest.

### Summary of Updates

all sections revised language, abstract reduced words, reference corrected, acknowledgement added, author list updated, minor update on all figures and captions

